# Modeling plant-microbe interactions with species distribution models

**DOI:** 10.1101/2024.07.12.603317

**Authors:** Zihui Wang, Sarah Piché-Choquette, Jocelyn Lauzon, Sarah Ishak, Steven W. Kembel

**Author notes:** **Corresponding authors** Zihui Wang Email address Steven W. Kembel.

## Abstract

Plants interact with diverse microorganisms that play a crucial role in plant growth and development. The diversity and distribution of plant microbiota are altered by anthropogenic environmental change, leading to subsequent impacts on ecosystems. Modeling the distribution of plant-associated microbes is critical for predicting and managing future changes in microbial function, but there remain challenges and open questions when developing these models. We present a conceptual framework for process-oriented predictive modeling of the distribution of plant-associated microbiota. We first describe different approaches to incorporating host plants into modeling microbial distributions, namely by including them as static variables, nesting them within microbial distribution models or incorporating them simultaneously via joint species distribution models. Additionally, we discuss issues associated with collecting and analyzing sequencing-based microbial data, emphasizing the importance of data normalization and careful interpretation of species distribution models. We further discuss how to incorporate evolutionary history into microbial distribution modeling. Finally, we present a case study demonstrating how incorporating host information can improve the prediction of microbial distributions. This study provides insights for predicting future distributions of plant-associated microbes under climate change and plant species redistribution, which can be generalized to other host-associated microbial systems.

## 1. Introduction

Plants interact intimately with diverse microorganisms including bacteria, fungi, archaea, protists, and viruses that inhabit both external and internal plant tissues (Berendsen et al., 2012; Fitzpatrick et al., 2020). These plant-microbe interactions comprise a range of symbiotic relationships including parasitism, commensalism, competition and mutualism, playing a pivotal role in shaping plant health and development (Pattnaik et al., 2021; Vandenkoornhuyse et al., 2015). While certain microbial groups such as mycorrhizal fungi, phytopathogens and nitrogen-fixing bacteria have been extensively studied over the last century, it was only in recent decades that the advent of high-throughput sequencing techniques revealed the remarkable and complex taxonomic and functional diversity of the plant-associated microbiota. Over the past decades, there has been a growing consensus on the importance of plant-associated microbiota for plant diversity, productivity, biogeography, and indeed nearly every aspect of plant ecology and evolution (Dastogeer et al., 2020; Delavaux et al., 2019; Hawkes et al., 2020). Moreover, plant-associated microbiota impact socio-economically important ecosystems such as agroecosystems and forests that are directly related to human well-being (Fisher et al., 2012; Thirkell et al., 2017). In this context, understanding the drivers of plant-microbe associations and elucidating the factors determining the diversity and distribution of the plant-associated microbiota have been considered key questions in ecology and evolution, as well as in plant sciences, forestry and agronomy.

Anthropogenic environmental changes — including climate change, nutrient deposition, land-use change and the introduction of invasive species — influence the diversity and distribution of plant microbiota both directly (i.e. by altering temperature) and indirectly (i.e. through altering plant traits or host species distributions) (Trivedi et al., 2022). These impacts are expected to prompt a rapid shift in the composition and function of plant-associated microbiota, given the shorter generation time and higher sensitivity of microbes to stresses compared to plants and animals (Cavicchioli et al., 2019). The alteration of the plant microbiota induced by global changes, on the one hand, represents a potential catastrophic risk to ecosystem health and human well-being. For example, the increased impact of phytopathogens under climate change has the potential to endanger food supplies for 10-60% of the world’s population and cause billions of dollars of economic damage annually (Bebber et al., 2014; Fisher et al., 2012). On the other hand, engineering the plant microbiota represents a potential approach to mitigate the negative effects of climate change on plants, for instance by enhancing plant resistance and acclimatation to abiotic and biotic stresses (Angulo et al., 2022).

Despite the critical role of the plant microbiota in mediating the effect of global change on plants and ecosystems — either accelerating or mitigating depending on the context and the microorganisms that are present — they have seldom been the focus of climate change studies and are almost always overlooked in policy development. A major challenge that prevents us from harnessing the full power of plant-associated microbiota is our shallow understanding of the response of microbes to environmental change (Cavicchioli et al., 2019). Therefore, there is an urgent need to develop process-oriented understanding and predictive modeling of plant-associated microbial distribution at large geographic scales. While recent studies have initiated investigations into the global diversity and biogeography of plant microbiota, i.e. the distribution of community diversity and composition (Li et al., 2023; Linde et al., 2018; Z. Wang et al., 2023), we are lacking a general framework to predict the geographic distribution of individual plant-associated microbial taxa.

This article presents a conceptual framework to address the challenges associated with developing process-oriented predictive models for the distribution of plant-associated microbiota. We begin by reviewing the use of species distribution models (SDMs) and their extensions, such as joint SDMs, for free-living microbes and host-associated microbes such as parasites. We argue that although these SDMs have offered insights into modeling the distribution of host-associated microbes, they are not readily applicable to plant microbiota due to 1) distinct constraints imposed on free-living and plant-associated microbes, 2) varying degrees of host specificity between the plant microbiota and parasites, and 3) differences in data types for plants and microbes. We then identify the major limitations of current SDM frameworks in modeling plant-associated microbiota including 1) challenges in integrating host-related biotic factors into microbial SDMs, 2) difficulties in handling microbial sequencing data, and 3) the evolutionary potential of microbes. To overcome these limitations, we propose different strategies for incorporating host information into microbial SDMs. These range from simply incorporating host taxa as static ‘environmental’ variables into microbial SDMs, to increasingly complex models such as nested SDMs and joint SDMs of plant-microbial interactions, which are further demonstrated using a case study on plant leaf-associated bacteria. Additionally, we discuss potential solutions for the compositional and highly dimensional nature of microbial sequencing data, as well as the hierarchical data structure and the challenges in collecting standardized microbial datasets. Lastly, we extend our discussion to the potential influence of microbial niche evolution on SDMs, proposing possible avenues for further exploration. Overall, the aim of this article is to initiate new discussions on SDM-based frameworks for modeling the distribution of plant-associated microbiota.

## 2. Recent progress and remaining challenges for modeling the distribution of plant-associated microbes

### 2.1 Species distribution models for microorganisms and parasites

While SDMs have been widely used to predict the distribution of plant and animal species across space and time (Elith & Leathwick, 2009), the characterization of microbial SDMs has historically lagged behind due to challenges in collecting microbial occurrence data, identifying microbial taxonomy and gathering environmental data at appropriate spatial and temporal scales (Hao et al., 2020). Consequently, microbial SDM studies have primarily focused on a few disease-causing microorganisms such as the human pathogen *Coccidioides immitis* and the plant pathogen *Pseudomonas syringae* (Gorris et al., 2019; Wang et al., 2018). Recent advances in environmental DNA approaches have facilitated the collection of microbial biodiversity data, opening new opportunities for modeling the distribution of microorganisms. For example, a recent study adapted SDMs to model the relative abundance of soil bacteria as a function of climate and edaphic variables, revealing that soil pH and climate can explain and predict the abundance distributions of most soil bacterial taxa (Mod et al., 2021). Another study focused on arbuscular mycorrhizal fungi, quantifying their distribution using climate and edaphic variables at a larger spatial scale, which provided a basis for predicting the geographic range shift of arbuscular mycorrhizal fungi under climate change (Davison et al., 2021). However, in the latter case of SDMs for plant-associated mycorrhizal fungi, a major limitation lies in the lack of consideration of host-related factors and the potential confounding effect between hosts and the abiotic environment (Kivlin et al., 2021).

The importance of host-related factors in shaping the distribution of host-associated microbes has been particularly emphasized in the context of parasitic interactions (Cunha et al., 2018; Gougherty & Davies, 2022; Kuhn et al., 2016; Piwowarczyk & Kolanowska, 2023). Studies on the SDMs of parasites have demonstrated that incorporating both host distribution and abiotic factors achieved higher accuracy in predicting the distribution of parasites, compared to considering only one of the two elements (Cunha et al., 2018; McDonough & Holloway, 2021). Depending on the data availability and the context of host-parasite interactions, host distributions can be incorporated into SDMs in various forms such as the presence-absence or abundance of host taxa, and the taxonomic, phylogenetic and trait attributes of host taxa and surrounding plant communities (Gougherty & Davies, 2022; Wisz et al., 2013). However, incorporating host distributions in SDMs also means the need for constructing additional SDMs to forecast future host distributions in order to predict the future distribution of parasites (Morales-Castilla et al., 2021).

Although SDMs for parasites have been refined to incorporate host distributions, they are not readily applicable to plant-associated microbiota. Several unique features of plant-associated microbial data pose challenges in modeling their distributions. First, plant-associated microbes differ from parasites in host specificity. Unlike many parasites which infect one or a few host species (though some parasites can have a wide host breadth, see (Park et al., 2018)), most plant-associated microbes live across a broad range of host plants and may also be found in non-plant habitats (Cordovez et al., 2019; Lajoie & Kembel, 2021). This makes it challenging to include the presence-absence of individual host species in the model. Secondly, microbial data are typically obtained through DNA sequencing, which provides relative abundances of microbial taxa such as amplicon sequence variants (ASVs) or operational taxonomic units (OTUs). This type of data raises additional challenges for microbial SDMs compared to parasites whose occurrence is generally measured through direct observation. For example, the presence-absence of microbes is highly sensitive to sequencing depth (Gloor et al., 2017) and synthesizing data from different studies is challenging since microbial taxa are often defined differently across studies (Brunel, 2020). Lastly, plant-associated microbial data are mostly collected from host individuals, leading to a hierarchical structure in microbial datasets where samples are not independent but clustered within host taxa or other host-related factors. Current parasite SDMs do not accommodate this hierarchical data structure (Björk et al., 2018).

### 2.2 Joint species distribution models and their application to host-associated microbes

Recent developments in joint species distribution models (JSDMs) present an opportunity for incorporating host information into modeling the distribution of plant-associated microbes. JSDMs extend traditional SDMs by simultaneously modeling the distribution of multiple species based on species attributes and environmental variables (Harris, 2015; Warton et al., 2015). JSDMs offer several advances over traditional SDMs. Firstly, JSDMs can hierarchically model distributions by incorporating factors that represent inherent relationships among species, such as species traits and phylogenetic relationships (Jamil et al., 2013; Ovaskainen & Soininen, 2011; Pollock et al., 2012). This allows JSDMs to consider that species with similar traits or phylogeny respond in the same way to the environment, and thereby improve predictions of distributions for rare and under-sampled species by using information from more abundant species that are phylogenetically close or have similar functional traits (Ovaskainen & Soininen, 2011). Secondly, JSDMs can infer associations among species based on model residuals, which reflect species co-occurrence patterns after controlling for the effect of environmental variables (Pollock et al., 2014). While the interpretation of this co-occurrence matrix is challenging because it can result from various processes such as missing predictors, species interactions and model misspecification, it provides information on species associations that can be used to improve predictions (Ovaskainen et al., 2017; Poggiato et al., 2021; Warton et al., 2015). Lastly, JSDMs can handle complex and hierarchical data structures by incorporating a variety of random factors such as sampling site, time, spatial distances, and other hierarchical layers (Björk et al., 2018; Ovaskainen et al., 2017). This improves the estimation of independent effects of factors and can provide information about the relative importance of different factors and processes.

Several recent studies have adapted JSDMs to predict the species distribution of host-associated microbes (Aivelo et al., 2019; Björk et al., 2018; Minard et al., 2019; Odriozola et al., 2021; Wang et al., 2023). For example, Björk et al. (2018) introduced a novel extension of JSDMs to accommodate the hierarchical structure of host-associated microbial data, which incorporates host-related factors as random terms in the model. This methodology makes it possible to discern the effects of host filtering among other ecological processes, as the authors illustrated with bird and sponge microbiota datasets. In another study, Wang et al. (2023) utilized JSDMs to characterize the geographic distribution of plant phyllosphere bacteria. They predicted the presence and abundance of keystone bacterial taxa as a function of host plant traits and climate. Their models incorporated data on the spatial arrangement and host phylogenetic relationships among samples as random terms to control for the effects of spatially and phylogenetically clustered data structure. This study revealed strong effects of host, climate, and spatial factors in controlling the occurrence and abundance of phyllosphere bacteria, suggesting that host-related factors and spatial proximity should be considered in modeling the distribution of the leaf-associated microbiota. Other studies that applied JSDMs to host-associated microbes reveal the potential of this approach in disentangling the importance of different processes, obtaining species co-occurrence patterns, and testing for niche conservatism along phylogenetic and trait axes (Aivelo et al., 2019; Minard et al., 2019; Odriozola et al., 2021). For example, Aivelo et al. (2019) used JSDMs to analyze *Ixodes ricinus* ticks’ bacterial communities along ecological gradients in the Swiss Alps, while Minard et al. (2019) studied the intraspecific variation of the gut microbial communities of *Melitaea cinxia* caterpillars to identify ecological factors driving their structure. Odriozola et al. (2021) used JSDMs to test whether shared habitat use or fungal-bacterial interactions were responsible for the co-occurrence observed between fungal and bacterial communities from deadwood.

### 2.3 Challenges of applying SDMs to plant-associated microbiota

In reviewing recent studies on SDMs for microbes and host-associated parasites, we have identified three categories of challenges of applying SDMs to model the geographic distributions of plant-associated microbes. The first challenge is related to incorporating host information into microbial SDMs, including what types of host data should be considered, how to deal with varying degrees of host specificity, and whether host distributions should be treated as a constant or dynamic data layer. The second challenge arises from the distinctive format and structure of microbial distribution data compared to plant and animal distribution data for which SDMs were initially developed. Microbial distribution data often come from field surveys of microbial communities based on DNA sequences, which are highly dimensional, compositional and can possess hierarchical structures. The last challenge is associated with the rapid evolution of microbes, leading to the possibility for rapid niche adaptation (Sousa et al., 2023) and host shifts (Vries et al., 2020) that should be considered in species distribution modeling.

## 3. Developing process-oriented predictive models for the plant-associated microbial distribution

### 3.1 Incorporating host information in SDMs of plant-microbe associations

To address the first challenge, we propose three scenarios for incorporating host plant distributions into the SDMs of plant-associated microbes. This is essential since microbial persistence requires host presence and a failure to incorporate information on host distributions may lead to a misestimation of the geographic range of plant-associated microbes. The first scenario is to incorporate plant distributions as a static layer that constrains microbial distributions. Secondly, the abiotic environment may influence the distribution of host plants, and thereby indirectly influence the distribution of plant-associated microbes. To account for this situation, plant species distributions can be incorporated into microbial SDMs as a dynamic layer, where plant distributions are modeled with a separate SDM and then used to predict microbial SDMs. The third scenario considers the feedback effect between plants and microbes, whereby plant distributions influence microbial distributions and vice versa. As a result, the distribution of microbes and their host plants may respond to environmental change simultaneously, which can be modeled using JSDMs by incorporating the co-occurrence patterns between plant and microbes (Fig. 1). Below we explicitly discuss these three types of SDMs in terms of rationale, data requirements, model structure and computational tools.

**Fig. 1.**
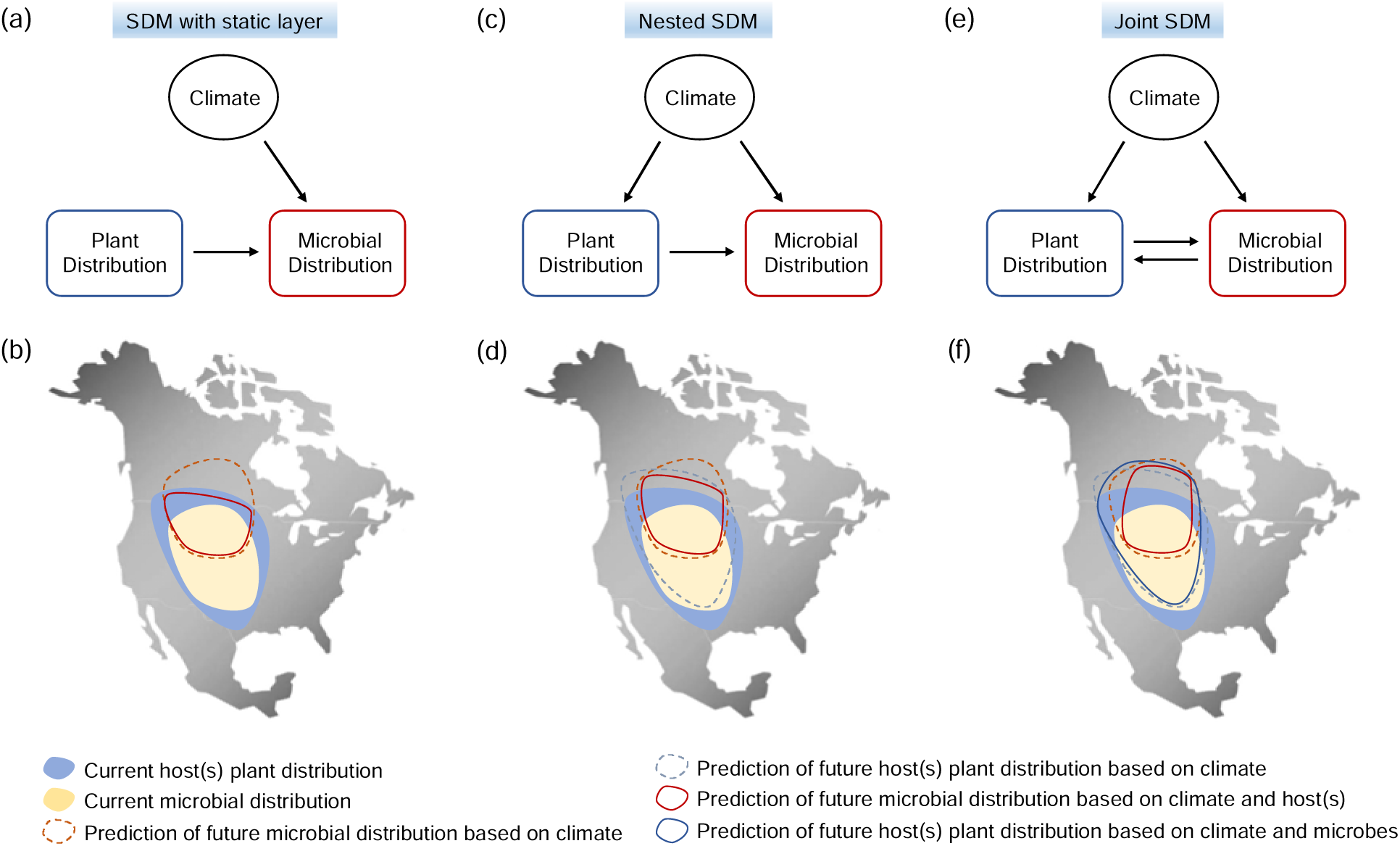
Conceptual frameworks for incorporating plant species distributions into models of plant-associated microbial distribution. Three scenarios describing the influence of plant distributions on microbial distributions are described, including (a,b) plant distributions as a constant constraint on microbial distributions, (c, d) environments affecting both plant and microbial distributions, with plant distributions additionally affecting microbial distributions, and (e, f) plant species and their associated microbes responding to environmental change simultaneously. (a, c, e) graphically summarize the modeling structures and (b, d, f) demonstrate the projection of current and future distributions of plant and microbial species under three different scenarios.

#### Scenario 1 ***–*** Incorporate host plants as a static layer

Host information can simply be included as a variable in the SDMs of plant-associated microbes. However, the way host information is represented in the model may depend on microbial data sampling strategies. In general, plant-associated microbial samples are collected from host individuals (“host-explicit”). Still, there are cases where microbial communities are sampled from a location with mixed hosts (“host-inexplicit”, e.g. Davison et al., 2021; Li et al., 2023). Although both types of microbial data can be used to construct microbial SDMs, they address different questions and require different model structures. Microbial SDMs based on host-explicit data predict the occurrence of microbial taxa on a specific host taxon across environmental gradients, which requires a hierarchical structure in the model to account for host effects. For this type of model, microbial occurrence in a location can be inferred by combining the occurrence of the host and the known associations between microbes and the host taxon. In contrast, models based on host-inexplicit data can directly predict the occurrence of microbes in a location as a function of biotic and abiotic environments, which is similar to plant and animal SDMs. Here, we primarily focus on microbial SDMs with host-explicit data although host-inexplicit SDMs are briefly covered.

Apart from microbial occurrence, data describing the host species, abiotic and biotic environments of the samples are required for plant-associated microbial SDMs (Fig. 2). Host information includes the species identity, traits, taxonomy, and phylogeny of the host individuals where the microbes are sampled (for host-explicit microbial data only). Biotic variables consist of information on plant communities where the microbes are situated, such as the presence-absence or the abundance of host plant species, as well as the taxonomic diversity, traits and phylogenetic properties of plant communities. The abiotic environment can include climate as well as microhabitat characteristics such as soil properties (in particular for root-associated microbes; Davison et al., 2021; Linde et al., 2018) and light conditions (in particular for leaf-associated microbes; see Carvalho & Castillo, 2018). Additionally, data on microbial attributes including taxonomy, phylogeny and functional traits can be incorporated into the SDMs (Fig. 2).

**Fig. 2.**
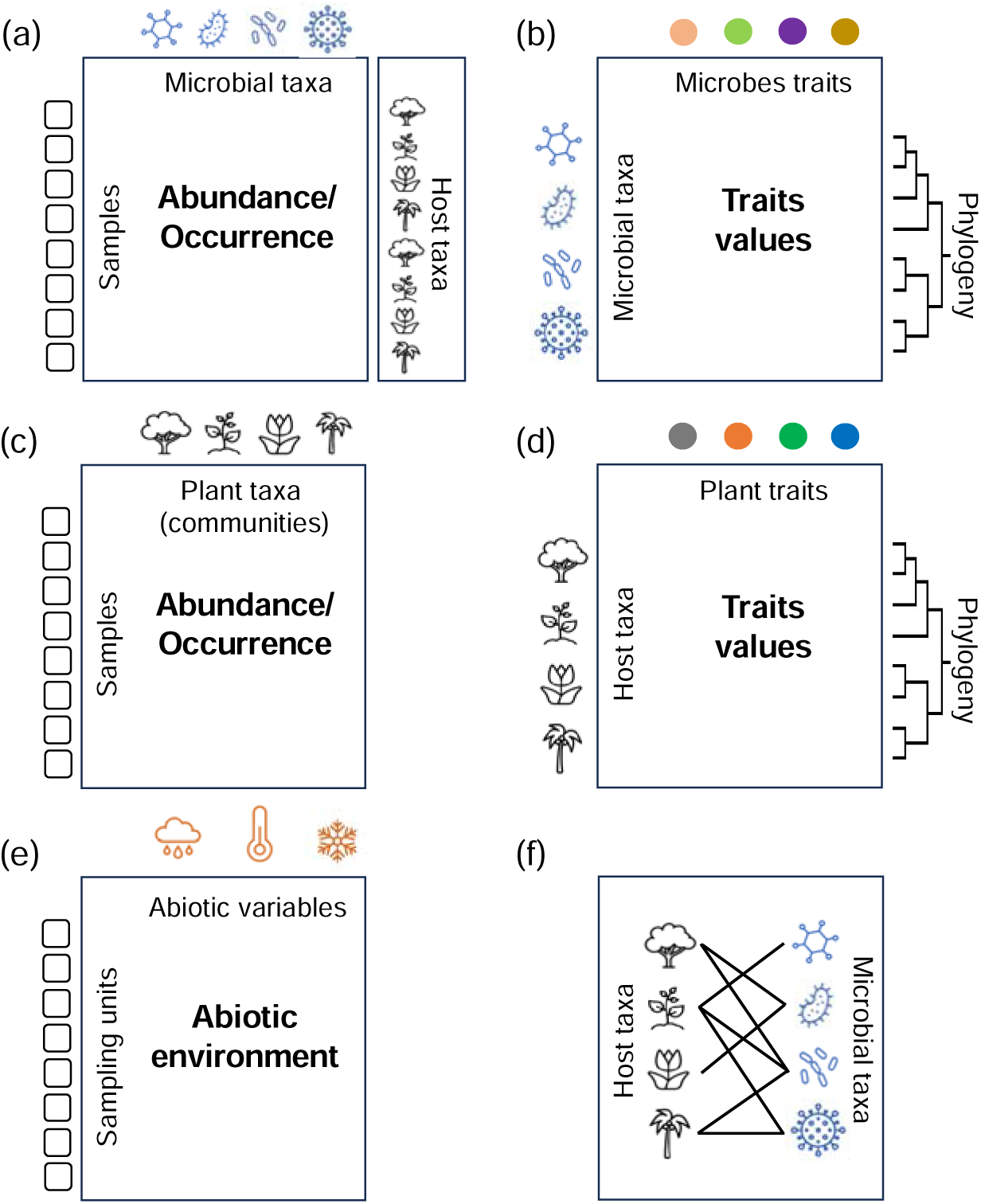
Data typically collected for plant-associated microbes. (a) Occurrence data of microbial taxa in each sampling unit. Sampling units are associated with host individuals and locations in host-explicit datasets, and only locations in host-inexplicit dataset; (b) Trait and phylogenetic data of microbial taxa; (c) Plant communities in which samples are collected; (d) Trait and phylogenetic data of plant species; (e) Abiotic environmental variables such as climate and edaphic variables, and (f) Plant-microbe interaction networks.

Below we illustrate how these datasets can be used to model the distribution of plant-associated microbes and generate predictions based on host plant distributions and abiotic environments (Fig. 1a). We start by modeling the presence or absence (PA) of plant-associated microbial taxon *j* in sample *i*. The subscript (*h*) indicates the host plant taxon from which sample *i* was collected:

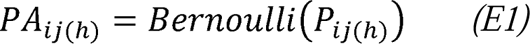

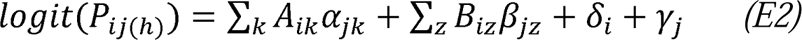

We use a probit regression to model the probability of presence P*_ij_*_(*h*)_ as a function of linear predictors in both fixed and random forms (E2). ∑*_k_ A_ik_α_jk_* represents the fixed effect of abiotic environments, where *A_ik_* denotes the abiotic factor *k* measured for sample *i* (e.g. temperature, soil nutrients or light), and the regression parameter *a_jk_* represents the response of microbial taxon *j* to abiotic factor *k* (Fig. 3). Similarly, ∑*_Z_ B_iZ_β_jZ_* represents the fixed effect of biotic environments, where *B_iZ_* denotes the plant community-related factor *z* measured for sample *i* (e.g. the abundance and diversity of host species in the plant community), and *β_jZ_* is the response of microbial taxon *j* to factor *z* (Fig. 3). The *δ_i_* and *γ_j_* terms represent sample- and microbial taxon-level random effects, which account for the difference in sampling design and taxon abundance, respectively.

**Fig. 3.**
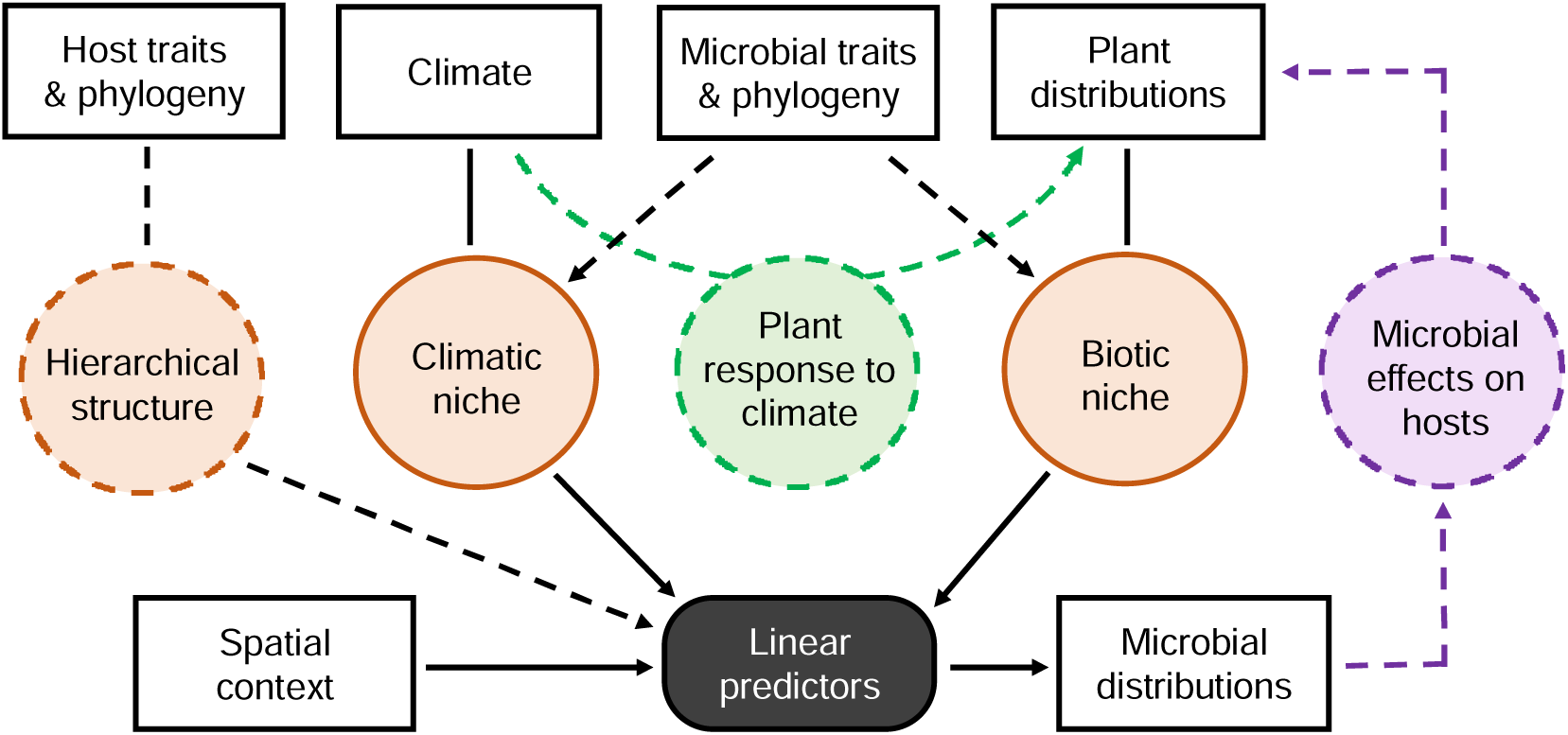
Graphical summary of the statistical framework integrating plant species distributions into microbial SDMs. Rectangles refer to data (explanatory or response variables), while circles represent parameters to be estimated. Plant distribution is integrated into the model as a static variable (orange color), dynamic variable (green) or as response variables in joint species distribution modeling (purple). Solid lines describe essential statistical inferences, while dotted lines refer to inferences that might be considered depending on the model structure and data availability (e.g. the hierarchical structure is included only when the microbial data is host-explicit).

The sample-level random term *δ_i_* provides an opportunity to accommodate the hierarchical structure of microbial sampling. Here we consider two types of hierarchy (Fig. 3). The first hierarchy arises from the nested microbial sampling within host plant species, where microbial communities collected from the same plant species are more compositionally similar compared to those collected from different plant species. To account for this effect, we included a random term *µ_h[i]_* to indicate the host species-specific effect on sample *i* (where *h*[*i*] is the plant species that sample *i* is collected from, see E3). This item can be further extended to include information on plant traits and phylogeny, which allows us to account for the observed trend that phylogenetically closer plant species tend to harbor a more similar composition of plant-associated microbes (see Björk et al., 2018 for detailed mathematics). The second hierarchy of microbial data is spatial hierarchy, in which microbial samples are nested within study sites, and microbial communities from the same site or nearby sites are more similar than those from spatially distant sites (Finkel et al., 2012). A random term *µ_s_*_[*i*]_ was included in E3 to indicate the effect of study site and spatial proximity (Ovaskainen et al., 2016).

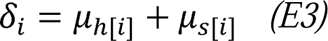

As explained before, SDMs based on host-explicit microbial data predict the occurrence of microbial taxa on a specific plant species. To determine the microbial occurrence at a location, we need to account for microbial occurrence on all plant species at that location, defined as:

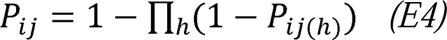

where *h* represents a given plant species at location *i*, and *P_ij(h)_* is the probability of presence of microbial taxon *j* on plant species *h* at location *i*.

In contrast, host-inexplicit microbial samples are collected from a location instead of a host individual. As a result, there is no need to consider host-specific random effects *µ_h[i]_* in the SDMs with host-inexplicit microbial data. In this case, the occurrence of microbes in a location can be directly predicted as a function of abiotic environments (such as climates) and biotic factors (including occurrence of potential host species or surrounding plant communities). However, such models are very sensitive to under-sampling (i.e. when not all plant species present at a location are sampled), in which case a random effect accounting for differential sampling effort should be considered (Morales-Castilla et al., 2021).

A variety of tools can be used for parameter estimation in the above models, including both single-species and multi-species distribution modeling. For example, generalized linear mixed-effect models (GLMMs) are often used to construct SDMs with hierarchical data, where factors such as host species identity and study site can be incorporated as random effects. With some extension, GLMMs can also be used to account for more complex hierarchical structures such as host phylogeny and spatial autocorrelation (Dormann et al., 2007). In addition, many multi-species distribution modeling tools such as JSDMs can model microbial SDMs while accounting for the random effects of host and space, which may take greater computing resource than single-species modeling because of the additional inference of the microbe-microbe co-occurrence matrix (Ovaskainen et al., 2017; Poggiato et al., 2021).

#### Scenario 2 ***–*** Incorporate host plants as a dynamic layer

Plant species distributions shift in response to environmental change, and therefore the future distributions of plant-associated microbes will depend on the redistribution of host species and the reassembly of plant communities. The model described above treats host plant occurrence as a static data layer; therefore, it can predict the geographic distributions of plant-associated microbes based on the current distributions of plant species. However, for predicting future microbial distributions, it is essential to construct separate SDMs to predict future plant distributions.

The model structure of plant SDMs has been established previously. Here we use a simplified plant SDM formula to illustrate how future plant distribution can be incorporated to predict microbial distribution. We model the presence-absence of host plant species *h* at location *i* (*PA_ih_*) using a probit regression, where the probability of presence (*P_ih_*) is modelled as a function of abiotic environments: ∑*_k_ A_ik_θ_hk_*, where the term *A_ik_* denotes the abiotic factor *k* measured for location *i*, and the regression parameter *θ_hk_* represents the response of plant species *h* to abiotic factor *k* (Fig. 3). These abiotic environments are not necessarily the same as those used in microbial SDMs since plants and microbes may respond to different environmental variables. *σ_i_* is the residual related to sample *i*, which can be extended to include other random effects if necessary.

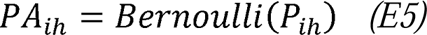

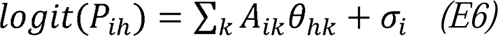

The reassembly of plant communities under environmental change can alter the values of plant community-related factors (*B_iZ_*) in E2, which represents the future properties of plant communities that the microbes may encounter. This value can be predicted in two different ways. One is through building plant SDMs for all possible plant species (E6) and extracting plant community information based on the predicted plant species distributions. The other is by building models to directly predict these factors, i.e. by modeling the abundance or diversity of plant species based on environmental variables. However, predicting plant community-level factors require additional data on plant community composition across environmental gradients (Fig. 3).

Based on the predicted occurrence of plant species *h* at location *i* (*P_ih_*) and the predicted occurrence of microbial taxon *j* on plant species *h* at location *i* (*P_ij_*_(*h*)_), the occurrence of microbial taxon *j* at location *i* can be calculated as E7, where *h* represents all potential host species of the microbial taxon *j*.

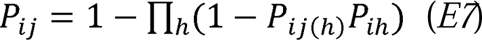

#### Scenario 3 ***–*** Predict plant distribution and microbial distribution simultaneously using JSDMs

While the models presented above focus on the influence of plant distributions and community properties on the distribution of plant-associated microbes, microbial distributions can have significant impacts on plant distributions. Many studies have emphasized the importance of plant-associated microbes in shaping plant species biogeographic distributions, for example, through positive effects of beneficial microbes on plant establishment under harsh environmental conditions, or negative effects of phytopathogens on plant fitness in otherwise suitable habitats (Angulo et al., 2022; Delavaux et al., 2019). Taking the feedback effects between plant and microbes into account when modeling the geographic distribution of plant-microbial interactions is essential for a realistic prediction of both plant and microbial SDMs.

The joint species distribution model (JSDM) approach, which has the feature of inferring and incorporating species associations into SDMs, provides a potential solution for accounting for the feedback effects between plant SDMs and microbial SDMs. To illustrate the use of JSDMs in predicting the distribution of plant-microbial interactions, we assume *k* microbial taxa and *n* host plant species collected at the same spatial scale. We predict the presence-absence of plant and microbes simultaneously by:

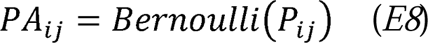

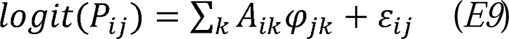

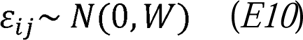

where *PA_ij_* is denoted as the presence of species *j* (*j* = 1…*k*…(*k*+*n*), including *k* microbial taxa and *n* plant species) at site *i*, which is modeled as a function of fixed term ∑*_k_ A_ik_φ_jk_* and a random term *ε_ij_*. The random term was drawn from a normal distribution with a mean of 0 and a variance-covariance matrix *W*. Here, *W* is designed to infer residual species-to-species relationships (hereafter residual correlation matrix), indicating the co-occurrence among species after controlling for the influence of fixed environmental terms. By inferring this residual correlation matrix in our model, we can estimate plant-microbe, plant-plant and microbe-microbe co-occurrence patterns. Although the inference of the residual correlation matrix in JSDMs does not necessarily change the estimation of fixed effects of environmental variables compared to single-species SDMs, it can alter the prediction of species distributions by incorporating information on species co-occurrence patterns as a random term (Poggiato et al., 2021). The interpretation of residual correlation is not clear, but it conveys biological information that JSDMs can utilize to better estimate species associations and to provide joint and conditional predictions (Wilkinson et al., 2021).

The conditional prediction of JSDMs based on residual correlation provides an opportunity to exploit the causality of the correlations between plant distributions and microbial distributions. For example, assuming that the observed plant-microbial co-occurrence is a result of plant species’ influence on microbial occurrence, one could build a JSDM to determine the probability of microbial occurrence given the occurrence of plant species (microbial occurrence conditional on plant occurrence). Similarly, JSDMs can predict plant occurrence conditional on microbial occurrence to test the influence of microbes on plant distributions. Additionally, plant occurrence and microbial occurrence can be predicted simultaneously as a conditional joint prediction (Fig. 3), assuming bidirectional effects between plants and microbes (Wilkinson et al., 2021). After building these different types of JSDMs, one could evaluate and compare the model performance to reveal insights into the causality of species co-occurrence between plants and their associated microbes (Wilkinson et al., 2021; Zurell et al., 2020).

### 3.2 Addressing challenges associated with microbial sequencing data

#### High dimensionality

The most commonly used method in collecting microbial distribution data is metabarcoding, a marker gene-based sequencing approach that amplifies a specific genomic region using short DNA strands as primers to detect sequences that are present and quantify their relative abundance. By comparing those amplified sequences to a database of reference sequences with known taxonomic identity, this method can simultaneously identify many microbial taxa within a sample. Microbial studies based on metabarcoding often cluster amplified sequences into operational taxonomic units (OTUs) based on their sequence similarity, or identify exact amplicon sequence variants (ASVs) from error-corrected sequences(Callahan et al., 2017). Both OTUs and ASVs can be used as the fundamental taxonomic unit in SDMs, serving as a pragmatic proxy for ‘species’ in microbial systems where traditional biological classifications are challenging. Metabarcoding of microbial communities often reveals a high microbial diversity, ranging from hundreds to thousands of taxa (Piché-Choquette et al., 2023). These highly dimensional community data pose a challenge given the computational cost of microbial JSDMs.

One potential way to reduce the high dimensionality of microbial community data is to focus either on a subset of microbial taxa of high research interest, or on a certain number of most abundant taxa, which discards rare species that are harder to model (Breiner et al., 2015). For example, keystone microbial taxa can exert considerable influence on microbial communities, and changes in the occurrence of these keystone taxa may cause drastic shifts in microbial composition and function(Banerjee et al., 2018). In this case, modeling the distribution of keystone taxa could be a priority. In addition, microbial SDMs may focus on microbial taxa with functions of interest such as nitrogen-fixing bacteria or mycorrhizal fungi. Focusing on a subset of microbial taxa can reduce the computational costs associated with JSDMs but requires prior knowledge of the keystone taxa in the study system or of the functional traits of microbial taxa. Another approach to reduce the high dimensionality of microbial community data is to model microbial distributions at a higher taxonomic level, such as the genus or family rank. For example, the bacterial genera *Pseudomonas* and *Methylobacterium* can be important for plant health (Mabood et al., 2014), and modeling the distribution of these plant-growth promoting bacterial genera is often of high research interest.

#### Compositional nature of microbial data

Metabarcoding-based microbial community data describe the relative abundance of microbial taxa. Most protocols for microbial metabarcoding attempt to normalize sequencing depth per sample to avoid introducing biases when comparing diversity metrics among samples, and as a result, variation in the number of sequences across samples do not reflect the variation in true abundance of microbes across those samples (Gloor et al., 2017). The compositional nature of metabarcoding-based microbial data poses two challenges for microbial SDMs. First, changes in the absolute abundance of a single taxon can alter the relative abundance of all other taxa in a sample, which may generate false correlations in the relative abundances among microbial taxa and between microbial abundances and environmental variables. Moreover, the total amount of sequences in a microbial community is influenced by microbial load (the number of microbial cells or amount of microbial DNA per sample), as well as DNA amplification protocols and sequencing library size (the number of DNA sequences obtained per sample). Microbial loads may vary depending on the unit volumes of microbial samples, and library size can be influenced by the efficiency of DNA extraction, PCR protocols, and other library preparation manipulations (McLaren et al., 2019). As a result, the total abundance of microbes in metabarcoding samples are, in general, biologically meaningless. That is, the total abundance of sequences does not necessarily reflect the true abundance of microbes in the ecosystem and the observed abundance of individual taxa is not comparable among samples. Therefore, it is inappropriate to use raw abundance data from metabarcoding to build microbial SDMs.

One solution to this issue is to estimate the absolute abundance of microbial taxa, for example, by combining metabarcoding with techniques that estimate total microbial abundance such as digital PCR and quantitative PCR (Barlow et al., 2020), or by inserting synthetic chimeric DNA spikes into microbial samples as a standard for absolute abundance estimation (Kim et al., 2021; Stämmler et al., 2016; Tkacz et al., 2018). However, methods for measuring absolute abundance are still in active development and are not as commonly used as relative abundance estimates in microbial studies (Morton et al., 2019). Relative abundance data of microbial communities can be potentially useful for microbial SDMs if sampling effort and library size are well controlled during data collection and analysis. For example, sampling effort can be standardized by collecting the same volume of material for all microbial samples, and library size can be normalized by rarefaction (Schloss, 2024). However, if the microbial loads and library sizes are not standardized during data collection, they must be considered during data analysis, potentially by incorporating sample volume and/or library size as a random term in SDMs (Odriozola et al., 2021). Overall, there is a lack of empirical data on the performance of these different approaches to normalizing abundances in microbial metabarcoding data, and studies are needed to determine the best approach to dealing with the compositional nature of microbial data in SDMs (but see Schloss, 2024).

#### Collection of large-scale microbial datasets

While the SDMs of plants and animals benefit greatly from the development of global observational datasets such as GBIF and eBird, collection of large-scale datasets for plant-associated microbes remains challenging. Numerous studies over the past decades have aimed to characterize microbial distribution at broad spatial scales, but most of them have focused on free-living microbes in habitats such as soils and oceans (e.g. Earth Microbiome Project; Thompson et al., 2017), or for a single host species (Human Microbiome Project Consortium; Methé et al., 2012) or specific microbial functional groups (GlobalFungi; Větrovský et al., 2019, 2020, and GlobalAMFungi; Větrovský et al., 2023). Large-scale studies on plant-associated microbes require not only a broad geographic range of sampling but also a wide host taxonomic coverage. This makes collecting data for host-associated microbes complex and logistically challenging.

While large-scale sampling campaigns to quantify plant-associated microbiota remain challenging, an alternative approach is to compile existing datasets to achieve a broad geographical and plant-host taxonomic coverage. An increasing number of datasets on plant-microbial associations has been published over the past decades (Sengupta et al., 2023). These data are accessible through international nucleotide sequence database collaborations such as the European Nucleotide Archive and the U.S. National Center for Biotechnology Information. These databases provide both sequence of the microbes and the metadata of samples such as the host plant, geographical location and the sampling protocol, representing a potential opportunity to investigate the global distribution and host specialization of plant-associated microbes. However, a significant obstacle for merging independent datasets together is the difference in the methods used across studies for sampling, DNA extraction, sequencing and data processing. These differences make it difficult to separate methodological artefacts from biological patterns (Brunel et al., 2020). Therefore, it is essential to control for technical biases and use standardized bioinformatics protocols to synthesize microbial datasets. Examples of this approach include a study that integrated sequences from different studies targeting different 16S rRNA gene regions by mapping them to near full-length 16S sequences based on a reference database (Lozupone et al., 2013). This reference mapping protocol provided a potential way to compare sequences from studies targeting heterogeneous regions of the 16S rRNA gene. Another study merged 30 independent bacterial datasets using a machine-learning approach that accounted for technical bias among studies (Ramirez et al., 2018). The authors showed that disparate amplicon sequences could be integrated to estimate bacterial community structure. However, there remain numerous challenges related to integrating microbial datasets across studies, and the development of standardized sampling, sequencing, and data analysis protocols will aid in the quantification of broad-scale plant-associated microbial distributions.

### 3.3 Incorporating microbial evolution into species distribution models

Microbes have the potential to evolve extremely rapidly given their high abundance, short generation time and the frequency of evolutionary events such as horizontal gene transfer and mutation. Consequently, microbes can quickly adapt to new environments and expand their geographic and host taxonomic range in response to climate change (Brennan & Logares, 2023; Chase et al., 2021). Incorporating this evolutionary potential into a SDM represents a key challenge for accurate prediction of microbial distributions. Many studies have demonstrated that the evolutionary history and functional traits of microbes were associated with their geographic and host range. For example, a study on the spatial distribution of soil bacteria showed that genomic traits and phylogenetic relatedness could be used to predict the habitat breadth of soil bacteria: bacteria with larger genomes and more metabolic versatility were more likely to have wider environmental and geographic distributions (Barberán et al., 2014). Another recent study on the host specialization of plant leaf-associated bacteria revealed that microbial genomic traits and phylogeny were associated with the host range of bacteria (Wang et al., in process). The importance of microbial phylogeny and traits in predicting their host range has also been broadly documented in the literature on plant-pathogen interactions, in which the potential hosts of poorly studied and newly identified pathogens were estimated based on host ranges of their relatives (Barwell et al., 2021; Gougherty & Davies, 2021; Kahn & Almeida, 2022; Marcot et al., 2023). Overall, these studies suggested that microbial traits and phylogeny provided valuable information of microbial niche evolution that can be used in models of the distribution of plant-associated microbes.

A potential way to incorporate microbial traits into SDMs is to assume that the microbial responses to abiotic and biotic factors (i.e. the *α_jk_* and *β_jz_* in E2) are not independent but correlated with microbial taxon-specific traits (Abrego et al., 2017). This can be achieved by including a nested model of *α_jk_* or *β_jz_*:

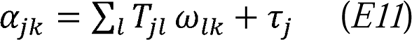

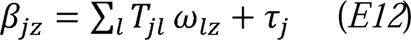

where *T_jl_* is the value of trait *l* for microbial taxon *j*, *ω_lk_* and *ω_lz_* represent the effect of trait l on the parameter *α_jk_* and *β_jz_*, respectively, and *τ_j_* is the error term. To account for the phylogenetic signal in microbial niches, the *τ_j_* should be drawn from a correlation matrix that assigns phylogenetically closer microbial taxa similar values, which can be indicated as

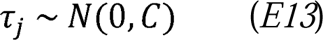

where *C* is a microbial taxon-taxon correlation matrix obtained from microbial phylogenies. However, by including *C* as the sole variance-covariance matrix, we assume that microbial niche is fully structured by the phylogeny, which is not always the case (Webb & Gaston, 2003). To acknowledge this, a parameter *ρ* is included to allow for varying strength of phylogenetic conservatism on microbial niche:

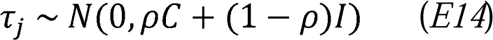

where *I* is a microbial taxon-taxon identity matrix and *ρ* ranges from 0 to 1, with a larger value indicating stronger effect of phylogenetic signal on microbial niche (Ives & Helmus, 2011).

## 4. Case study

We applied SDMs to a dataset of plant leaf-associated bacteria to demonstrate how the future distribution of plant-associated microbes can be predicted under different scenarios of climate change and host species redistribution. The dataset was released from a biogeographic survey of leaf microbes in 10 forest sites along a latitudinal gradient (18.7°N-47.1°N) in China (Wang et al., 2023). A total of 1453 microbial samples from 329 plant species were collected and sequenced using metabarcoding of the bacterial 16S rRNA gene to quantify bacterial community composition, revealing a total of 49,254 bacterial amplicon sequence variants (ASVs). To reduce the dimensionality of this dataset, we selected a small subset of the data by focusing on 12 of the most common plant species and 30 of the most abundant but not ubiquitous bacterial ASVs. The resulting dataset consists of 188 microbial samples nested within 12 plant species and 10 forest sites. The metadata of these samples included, apart from the site and plant species, three site-level climatic variables: annual mean temperature, annual precipitation, and temperature isothermality; and three host plant attributes: specific leaf area, leaf nitrogen concentration, and leaf phosphorus concentration. In addition, the presence-absence of host species in plant communities across study sites was included. Overall, this dataset represents a broad geographic and host species coverage that is essential for a large-scale SDM of host-associated microbes, though its spatial grain is limited by the relatively small number of study sites.

We built three models to predict the distribution of bacterial ASVs as a function of three climatic variables and three host attributes while taking the hierarchical structure of the data into account. Bacterial ASV community data (i.e. 188 samples and 30 ASVs) – the response variables – were transformed to presence-absence values before probit regression was applied. In the first model, we considered host plants as a static layer, where the 3 host plant leaf traits and the 3 climatic variables were incorporated as fixed effects (i.e. A_1_-A_3_ and B_1_-B_3_ in E2). The model included study site and plant species identity as two random terms to account for the variation in bacterial presence-absence within study sites and plant species (i.e. *δ_i_* in E2 and E3). In the second model, we considered host plants as a dynamic layer: the distribution of plant species was first modeled as a function of the 3 climatic variables following E6, where the presence-absence of 12 plant species across 10 study sites was used as the response variable in a probit regression. Following E7, we estimated the probability of presence for each bacterial ASV based on the predicted occurrence of host plant species. Finally, in the third model, we predicted plant and bacteria distributions simultaneously with joint species distribution modeling. The presence-absence of 30 bacterial ASVs and 12 plant species were combined as the response variables, 3 climatic variables were used as fixed effects, and study site was treated as a random effect. All these models were estimated using Markov Chain Monte Carlo (MCMC) implemented within the R package ‘Hmsc’ (v.3.0.13) assuming the default prior distributions (Tikhonov et al., 2020).

The first model, where host plants were considered a static variable, explained ∼57% of the variation in the presence-absence of leaf bacterial ASVs, most of which was explained by climatic variables (particularly annual mean temperature) (Fig. 4). We found that considering hosts as a dynamic layer expanded the estimated thermal range of leaf bacteria compared to models where host occurrences were treated as static variables (Fig. 5). This indicated that the range shift of host species can influence the prediction of bacterial distributions. However, predicting bacteria and plant distributions simultaneously with joint species distribution modeling only slightly altered the range of predicted bacterial distribution, where the range of some bacterial ASVs distributions expanded while others shrank (Fig. S1).

**Fig. 4.**
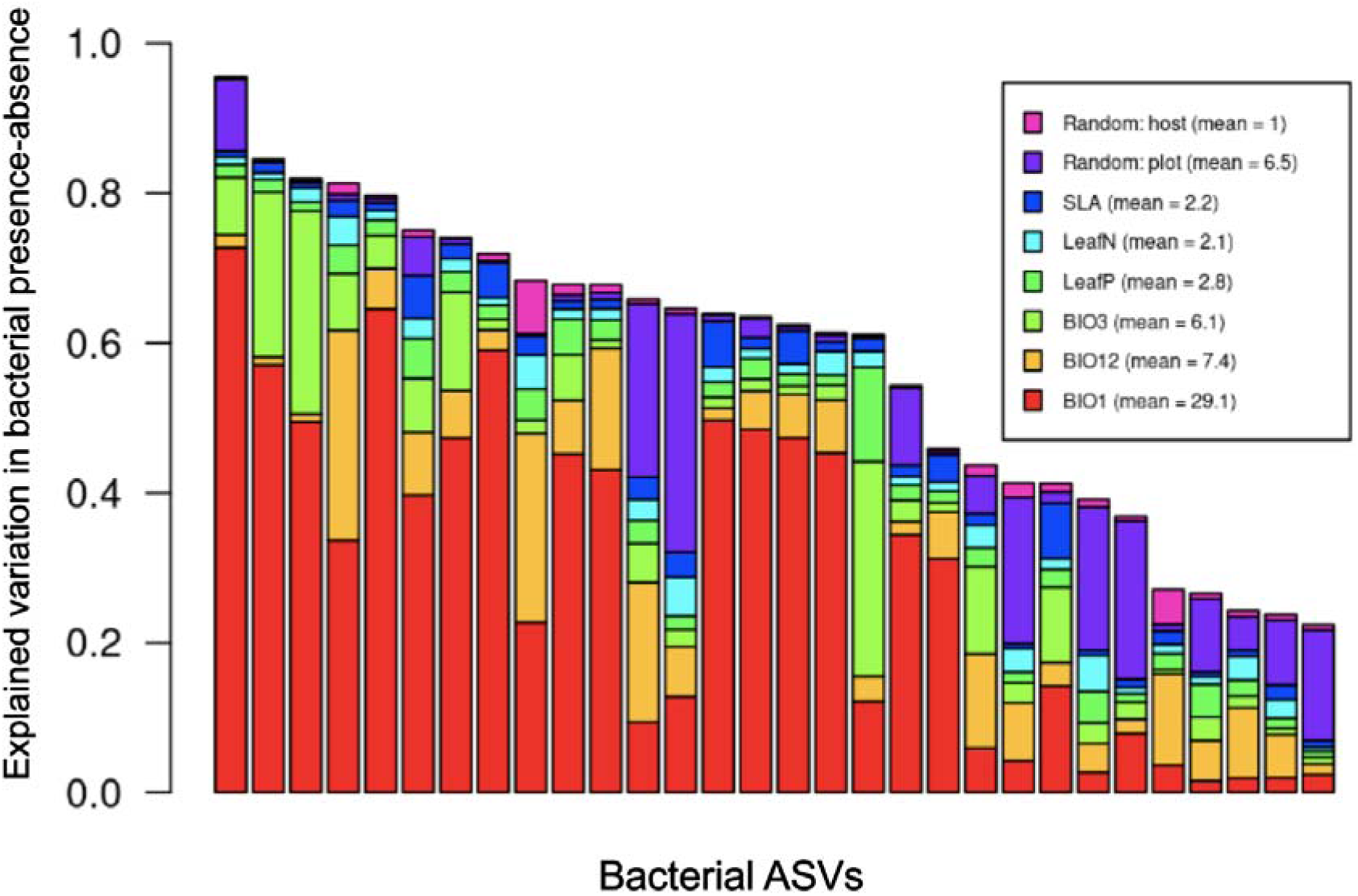
Explained variation in the presence-absence of bacterial ASVs in a species distribution model with host attributes considered as static variables. Each column represents a bacterial ASV, ranked in order of total explained variation. Colors represent the variation explained by each variable as noted in the caption box. Abbreviations: Random host and plot, the random effect of host plant species and study plot; SLA, specific leaf area; LeafN and LeafP, leaf nitrogen and phosphorus concentrations; BIO1, BIO12 and BIO3, the annual mean temperature, annual precipitation and temperature isothermality.

**Fig. 5.**
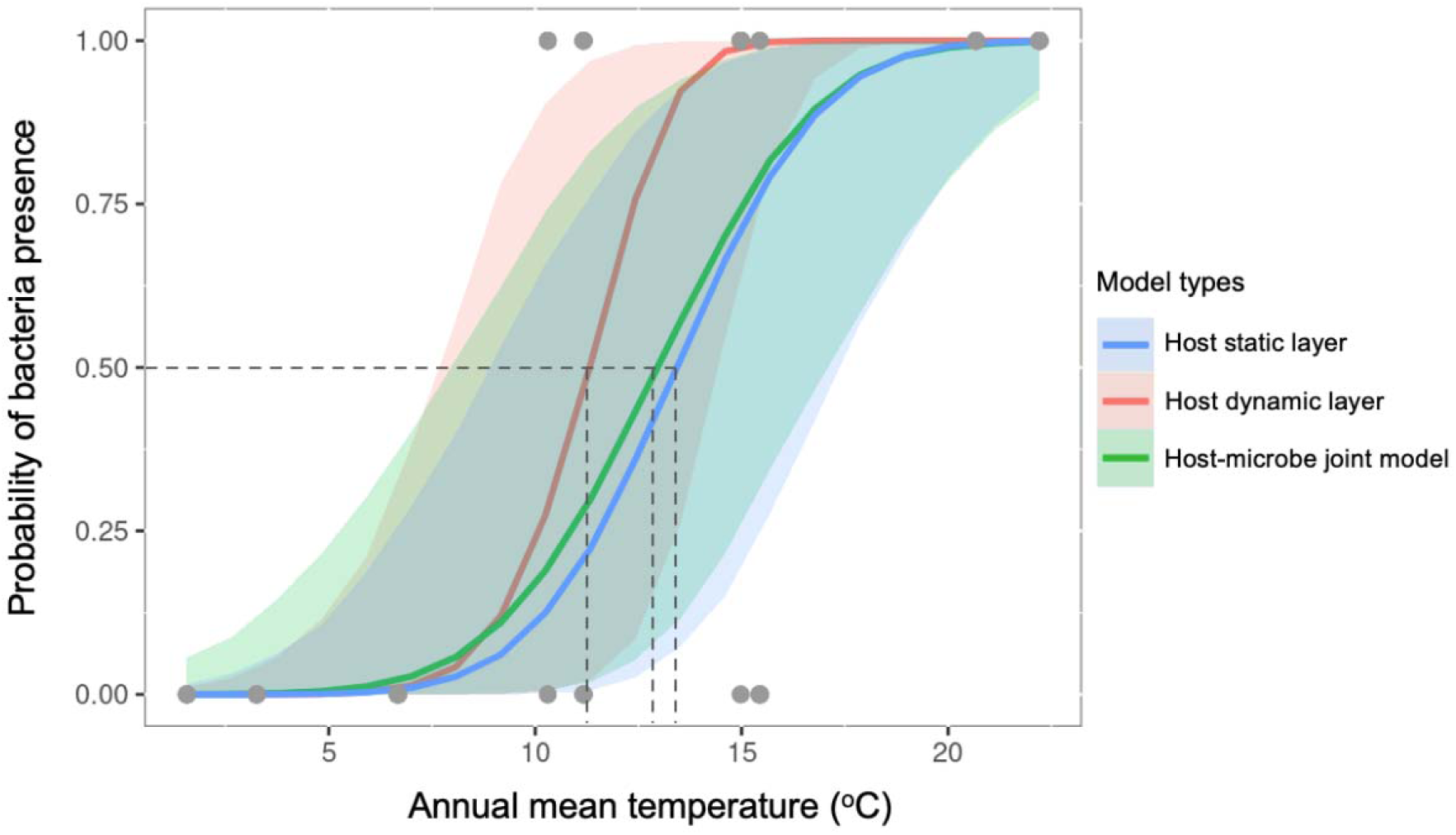
Predictions of bacterial presence across a temperature gradient based on three models, where host plants were considered as a static variable, a dynamic variable and a joint response (model type). The figure demonstrates the prediction for an example bacterial taxon (ASV_16); predictions for the other ASVs can be found in Supplementary Figure 1. The mean and 95% confidence intervals of the predictions are shown. Dashed lines indicate the temperature at which each model predicts a 50% probability of occurrence.

Overall, this case study suggests that incorporating the dynamics of host plant distributions is important for predicting the distributions of plant-associated microbes (Morales-Castilla et al., 2021). The persistence of plant-associated microbes relies on the presence of host plants, and therefore ignoring the range expansion of host plant species may underestimate the future distribution of plant-associated microbes, as illustrated in Fig.1. At the same time, the geographic distribution of plant species can be influenced by their associated microbes. For example, the lack of beneficial microbes in a new habitat may limit the expansion of plant species (Delavaux et al., 2019), while the lack of pathogenic microbes could facilitate plant invasion and expand plant species distribution (Keane & Crawley, 2002). This feedback effect between plant distributions and microbial distributions can be represented by a residual plant-microbe association matrix in the joint species distribution modeling framework, which is useful to predict the distribution of plant and microbes while considering the complex plant-microbial interactions (Poggiato et al., 2021; Wilkinson et al., 2021).

## 5. Future directions

Predicting how plant communities and their associated microbes will respond to environmental changes will be crucial in developing ecosystem management policies aiming to prevent or mitigate phytopathogen invasions and preserve ecosystem services provided by microbes to human populations. The conceptual framework presented in this review is an attempt to orient future research in the field of microbial biogeography by addressing key challenges and proposing alternatives to model the future distribution of host-associated microbes. Promising future directions for the study of host-associated microbial distributions include collecting large-scale datasets on microbial distributions, developing novel statistical and computational approaches to model microbial distributions, and assessing the validity of models of host-associated microbial distributions.

### Need for large-scale data and standardized protocols

An important future direction for modeling plant-associated microbial distributions will be the collection of large-scale standardized datasets to allow modeling of distributions at global scales. Given the difficulty of comparing microbial community data collected using different technologies and protocols (Kool et al., 2023), a collective effort should be made to encourage the use of consistent and documented protocols as a roadmap for collecting and analyzing environmental samples yielding plant-associated microbial data.

### Need for computationally efficient modeling approaches

A major bottleneck in the development of host-associated microbial SDMs is that building SDMs from large and complex datasets is computationally intensive. Strategies to reduce model complexity and computing costs when fitting microbial SDMs remain to be evaluated quantitatively. We suggest that a promising research priority will be to compare different approaches such as model selection, ridge regression, dimensionality reduction, or other statistical methods that reduce the complexity of host-associated microbial SDMs. Another way to reduce computational demand while slightly compromising the robustness of the model is to use less greedy species distribution modeling approaches, such as approximated Bayesian methods like Integrated Nested Laplace Approximation and Variational Bayes (Blei et al., 2017; Rue et al., 2009). Lastly, advances in computational power such as the use of GPU parallel computing should permit the analysis of increasingly complex models (Pichler & Hartig, 2021).

### Need to test and validate distribution models

Finally, an important future direction will be to assess the validity of predictive models of microbial distributions, by verifying and comparing true microbial occurrences in ecosystems against the predictions obtained with SDMs. For example, in the short term, this could be achieved by modeling the distribution of some chosen microbial taxa with data taken from a restricted spatial range, and collecting samples along a spatial gradient within and beyond their predicted range to survey their presence and test the accuracy of the SDMs. In the medium term, another possibility to test the predictions of SDMs would be to model the distribution of microbes associated with invasive plant species, since these plants tend to rapidly expand their geographic range under climate change (Dukes et al., 2009; Wang et al., 2019). By doing so, researchers might be able to compare in just a few years from now the true novel microbial distributions against their predicted distributions computed with presently available data. These kinds of assessments will also be an invaluable opportunity to compare the performance of competing model types, with the aim of gaining a better understanding of which approach is best suited for different study systems and data structures. Ultimately, our ability to better model the distributions of host-associated microbes will make it possible to predict potential responses of plants and their associated microbes to global change.

### Data accessibility

The data that support the findings of the case study are openly available in Sequence Read Archive at https://www.ncbi.nlm.nih.gov/sra/PRJNA1001021, with accession number: PRJNA1001021

